# Nitrite deteriorates bioreactor performance by reducing growth of *Ca.* Brocadia sapporoensis instead of inhibiting the anammox activity

**DOI:** 10.1101/2023.11.24.568540

**Authors:** Xuejiao Qiao, Liyu Zhang, Yang Wu, Chunfang Deng, Yichi Zhang, Xue Zhang, Yan Yan, Weiqin Zhuang, Ke Yu

## Abstract

Effects of nitrite on anammox activities have been of widespread concern. However, the molecular mechanisms of specific microorganisms in anammox systems responding to nitrite remain unexplored. This study investigated the effects of excess nitrite on core metabolism of AnAOB and symbiotic bacteria, and further elucidated the response mechanism of these effects on microbial growth and nitrogen removal performance. Specifically, nitrogen removal process in a continuous-flow anaerobic ammonia oxidation membrane bioreactor completely collapsed when the nitrite concentration reached 243 mg N/L. Integrated meta-omics analyses demonstrated that excess nitrite disrupted the energy metabolism of *Ca.* Brocadia sapporoensis (AMXB1), reducing the energy available for establishing tolerance. It disrupted cell replication by impairing biosynthesis process of AMXB1, especially DNA replication and the formation of vital cell structures, e.g., cell membrane and cell wall, as well as the cellular protection system, leading to the collapse of the anammox system. In addition, the cross-feeding of glycogen, lipopolysaccharide and amino acid between AMXB1 and symbiotic bacteria was hindered by excess nitrite, which also contributed to the anomalous cell proliferation and metabolism of AMXB1. These findings contribute to our understanding of the ability of anammox consortia to respond to nitrite stress and process stability in engineered ecosystems.

**Highlights:**

- NO ^-^-N concentration of 243 mg N/L caused the performance collapse of a continuous-flow anammox MBR.
- Excess nitrite likely disrupted the energy metabolism of AMXB1, reducing the energy availability for mitigating nitrite toxicity.
- The cross-feeding between AMXB1 and symbiotic bacteria was hindered by excess nitrite.
- The hindrance of cross-feeding was reversed as the concentration of nitrite decreased.

**Graphical abstract:** 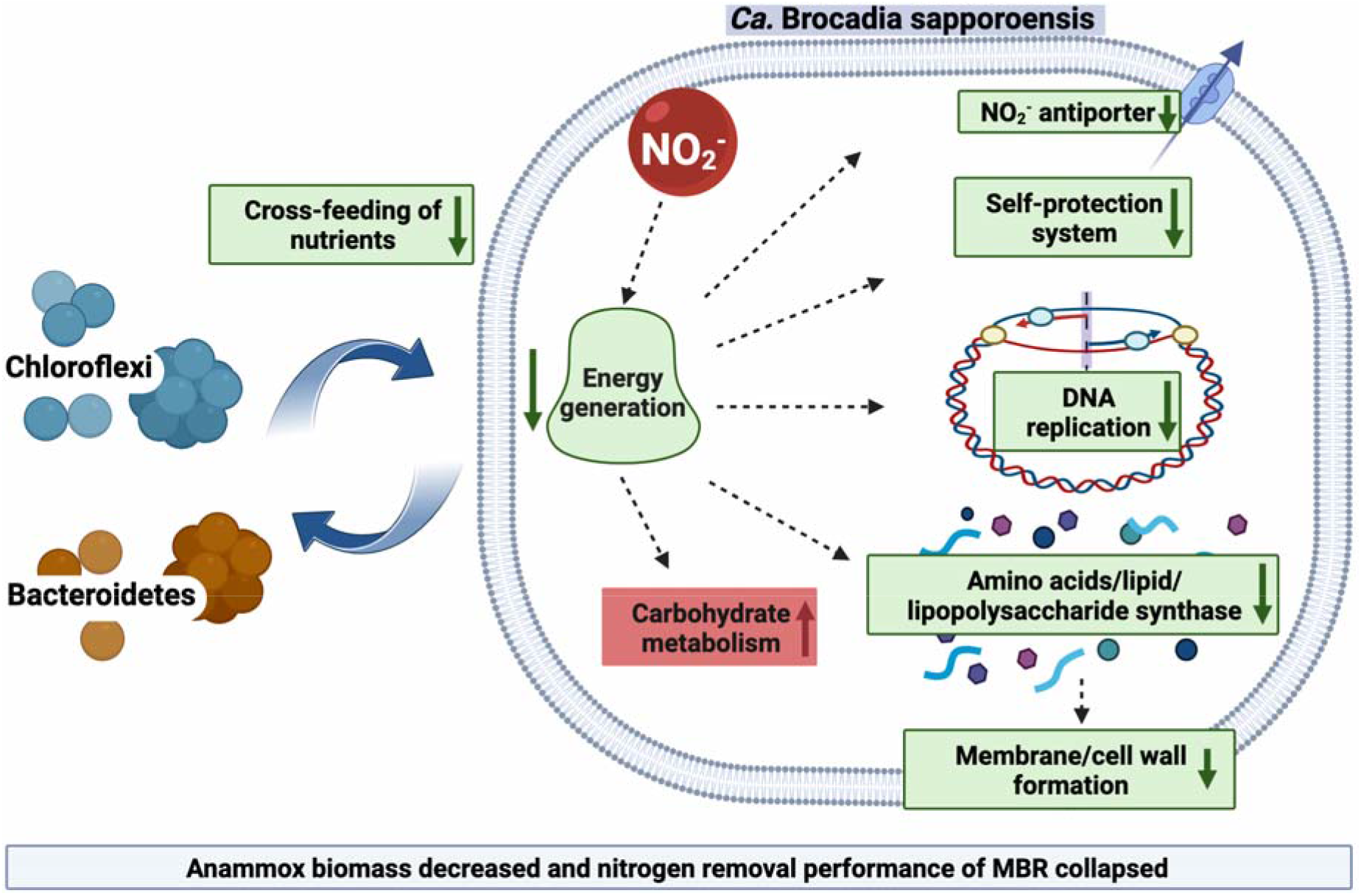

## 1. Introduction

As an efficient, economical and environmentally friendly biological nitrogen removal process, anaerobic ammonia oxidation (Anammox) has undergone significant development^1^. Anammox is mediated by the autotrophic anaerobic ammonia-oxidizing bacteria (AnAOB), which utilize nitrite (NO ^-^-N) as an electron acceptor to oxidize ammonium (NH_4_ ^+^-N), producing nitrogen gas (N_2_) and nitrate (NO_3_ ^-^-N), under anaerobic conditions ^2^. Anammox finds wide application in treating high-concentration ammonium (400 - 6000 mg/L) wastewater including landfill leachate and pharmaceutical, hospital, and livestock wastewater ^3-5^. Maintaining an optimal nitrite concentration is essential for stable nitrogen removal in the anammox process ^6^. When using anammox process to remove ammonia from wastewater, careful control is needed to oxidize roughly 50% of the ammonia load to nitrite without being further oxidized to nitrate, which has no use to AnAOB ^7,8^. Nitrite is a potent inhibitor and bactericide, even at low concentrations. The accumulation of nitrite during the anammox process can result in the complete inhibition of AnAOB and other bacteria, causing irreversible impact and long recovery.

Many studies have demonstrated the inhibitory effect of nitrite to AnAOB ^9^. However, the threshold concentrations of nitrite reported in the literature varied significantly, ranging from as low as 30 mg N/L to as high as 768 mg N/L ^10-14^. Notably, there are variations in the specific conditions used to determine the inhibitory nitrite concentrations in the aforementioned studies, making it challenging to draw a consensus reference concentration to avoid reactor collapse caused by excessive nitrite.

Another important feature of AnAOB, as a group of autotrophic bacteria, are always associated with some heterotrophic bacteria, establishing some mutual beneficial interactions. Among these interactions, cross-feeding might play a crucial role in the nutritional interaction between AnAOB and other symbiotic bacteria, facilitating exchanges of nitrogen, vitamins, cofactors, and amino acids ^18-20^. Autotrophs such as AnAOB can release soluble products and extracellular polymeric substances, which serve as a source of organic compounds for other symbiotic bacteria ^21-23^. Heterotrophs, including canonical denitrification (DN) and dissimilatory nitrate reduction to ammonium (DNRA) bacteria, can use these organic compounds as electron donors to drive dissimilatory nitrogen metabolisms ^24-26^. Additionally, certain bacterial taxa such as Armatimonadetes and Proteobacteria provide folate and other important cofactors for the carbon fixation by AnAOB ^18,19^. While members from the genus *Ca.* Brocadia can synthesize most amino acids, their growth might enhanced by threonine, methionine, and histidine secreted by some Proteobacteria ^20^. Therefore, a cross-feeding relationship is essential for efficient nitrogen removal in AnAOB consortia ^27^. The cross-feeding of nutrients within anammox systems has the potential to regulate and influence the activity of AnAOB ^18^. Yet, there has been no study investigating the mechanisms of cross-feeding within microbial communities during prolonged periods of excessive nitrite in anammox systems.

Consequently, while exploring the mechanisms behind nitrite inhibition on the anammox processes, some studies suggested that the inhibitory effect could be attributed to the nitrite’s inhibitory effect to both AnAOB and the symbiotic bateria with the ananmmox communities. Excess nitrite may promote the growth of other microorganisms and reduce the living space available for AnAOB without additional carbon source, thus compromising the nitrogen removal performance of the system ^17,28^. Additionally, some researchers proposed that the inhibition could be attributed to free HNO_2_ ^16^, particularly in acidic conditions. However, the exact inhibitory mechanism of nititrite inhibition has yet to be fully elucidated.

The objective of this study was to elucidate the transcriptomic response mechanism of AnAOB and the shift in the cross-feeding relationship within anammox consortia during the collapse of the anammox system induced by excessive nitrite. A long-term nitrite inhibition experiment was conducted in an anammox membrane bioreactor (MBR) dominated by *Ca.* Brocadia, with progressively increasing nitrite loadings until the collapse of the system. Subsequently, to restore the nitrogen removal performance of the MBR, the nitrogen loading rate (NLR) was reduced by half. Genome-resolved metagenomic and metatranscriptomic analyses were performed to investigate the impact of nitrite stress on nitrogen cycling, as well as the intensity of carbohydrate, amino acid, coenzyme, and vitamin synthesis in *Ca.* Brocadia and its symbiotic bacteria during both the collapse and recovery periods. The findings provide valuable insights into the mechanisms underlying bacterial growth and the nitrogen removal performance of the anammox system under these conditions. This research contributes to establishing a theoretical foundation and offering technical support for mitigating nitrite inhibition in the anammox process.

## 2. Materials and methods

### 2.1. The Membrane Bioreactor (MBR) operation

In this study, a continuous flow complete-mix MBR system with a 1.5 L effective volume was utilized to conduct the experiments. The detailed configuration and operations of the MBR were reported previously ^6^. In summary, seed sludge was obtained from a laboratory-scale anammox reactor that had been operating for two years at a nitrogen loading rate (NLR) of 2 g N/L/d. After inoculation, the initial concentration of mixed liquor volatile suspended solids (MLVSS) in the MBR was 0.8 g/L. The predominant AnAOB species was *Ca.* Brocadia with an abundance of about 50%. Synthetic wastewater medium was sparged with N_2_ for 20 min to remove dissolved oxygen before fed into the MBR continuously using a peristaltic pump, resulting in a hydraulic retention time (HRT) of 24 h. Mixing in the MBR was achieved by a mechanical stirrer operating at 150 rpm. Dissolved oxygen in the MBR was controlled at less than 1% air saturation. Additionally, the temperature was maintained at 35±1°C, while the pH ranged between 6.8 and 8.0.

### 2.2. Long-term nitrite inhibition

The anammox MBR was operated for a total of 205 d. This period can be divided into five stages, as shown in Table 1. In the proliferation period (days 1-43), the AnAOB-containing sludge graduately adapted to the new environment and the synthetic wastewater media. NLR was gradually increased once the nitrogen removal efficiency reached almost 90% and remained stable for two consecutive HRTs. In the stabilization period (days 43-60), influent NH_4_ ^+^-N and NO_2_ ^-^-N concentrations were all kept constantly at 1000 mg N/L to maintain a stable environment for the subsequent experiments. In the inhibition period (days 60-96), the influent NH_4_ ^+^-N concentration was maintained at 1000 mg N/L, while the influent NO_2_ ^-^-N concentration was increased from 1000 to 1100 mg N/L to induce nitrite stress in MBR. In the collapse period (days 96-135), the NO_2_ ^-^-N concentration was further increased to 1200 mg N/L until the anammox system experienced a complete collapse. In the recovery period (days 135-205), the nitrogen removal performance of the anammox process was restored by first decreasing half NLR to 1000 mg N/L/d and then gradually increasing it again to 2000 mg N/L/d.

**Table 1.**
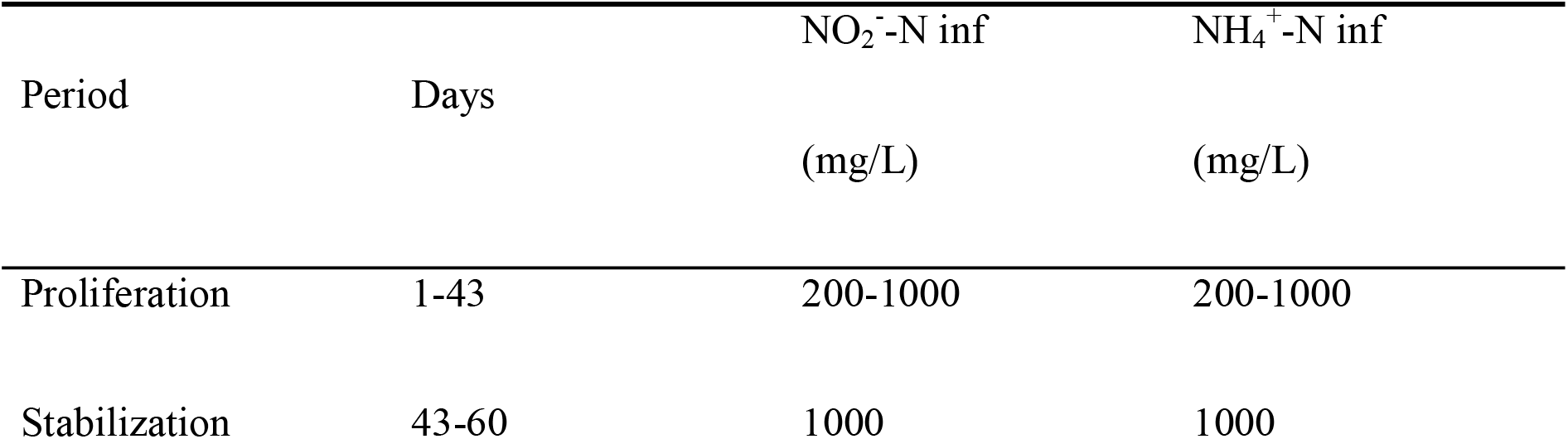
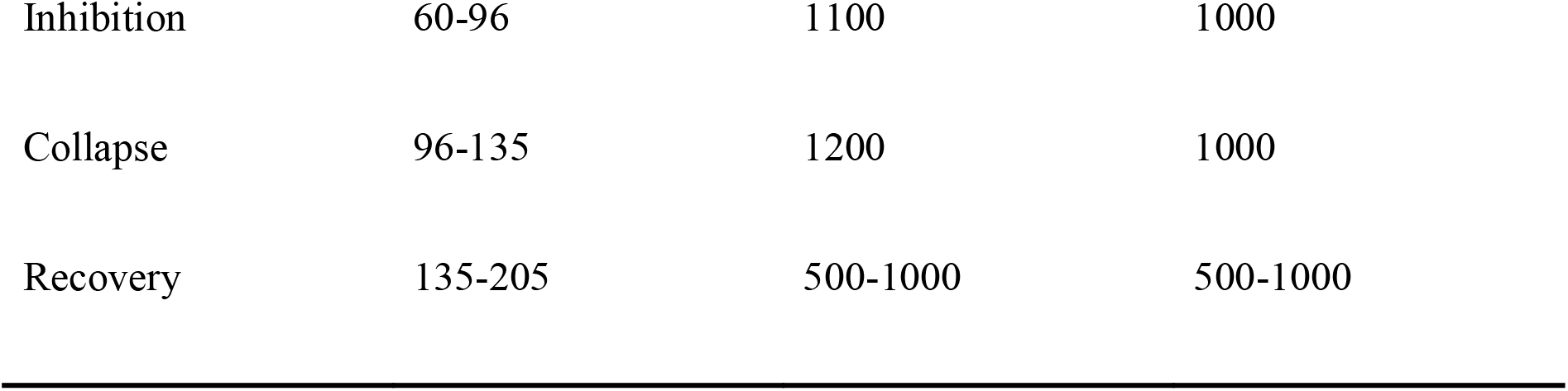
Experimental periods during reactor maintenance.

### 2.3. Water quality and sludge characteristics

Influent and effluent samples were collected for chemical analyses two or three times a week. Samples were filtered using 0.22-μm membrane filters (JIN TENG, China) before analyzing the concentrations of nitrite (NO_2_ ^-^-N), ammonium (NH_4_ ^-^-N), nitrate (NO_3_ ^-^-N) using the standard spectrophotometric method (APHA, 2012). Total organic carbon (TOC) in the effluent samples was measured using the standard combustion method (APHA, 2012). To measure MLVSS (biomass concentrations), four sludge samples in the stabilization, inhibition, collapse, and recovery periods were collected and measured using the Standard Method (APHA, 2005). Additionally, the total protein concentrations of the sludge was utilized as an additional indicator of biomass concentration. 5 mL mixed liquor was collected from MBR, centrifuged at 4000 rpm for 10 min, and then washed with Phosphate Buffered Saline (PBS, 0.1M and pH 7.5) for three times. A total of 21 sludge samples obtained throughout the entire operation period were subjected to protein extraction using the Total Protein Extraction Kit (3101, BestBio, China). The protein concentrations were determined using the Enhanced BCA Protein Assay Kit (P0010, Beyotime, China).

### 2.4. Sampling, extraction and sequencing of DNA and RNA

Sludge samples were collected at 22 time points during the experiment from MBR for DNA extraction using the DNeasy PowerSoil Pro Kit (QIAGEN, Germany). Sequencing of DNA followed the methods described in our previous study ^6^. In brief, the hypervariable V4 region of the bacterial 16S rRNA gene was amplified using primers 515F (5′-GTGCCAGCMGCCGCGGTAA-3′) and 806R (5′-GGACTACHVGGGTWTCTAAT-3′). The sequencing library was generated using NEBNext^®^ UltraTM II DNA Library Prep Kit (New England Biolabs, MA, United States) and sequenced on an Illumina Nova6000 platform generating 250-bp paired-end reads (Guangdong Magigene Biotechnology Co., Ltd., Guangzhou, China). For shotgun metagenomic sequencing, a subset of 19 DNA samples were selected in the five periods of the experiment. DNA quality was assessed prior to library preparation and sequenced on an Illumina NovaSeq 6000 platform generating 150-bp paired-end reads (Guangdong Magigene Biotechnology Co., Ltd., Guangzhou, China).

From the experiment, a total of 15 sludge samples were selected from five periods with significant changes in nitrogen removal performance for metatranscriptomic analyses: day 49, 56 and 63 from the stabilization period; day 70, 77 and 84 from the inhibition period; day 91, 98 and 105 from the start of collapse period; and day 120, 130 and 135 from the end of collapse period; day 184, 197 and 205 from the recovery period (Figure 1). Total RNA was extracted from these samples using the RNeasy PowerSoil Total RNA Kit (QIAGEN, Germany). rRNA removal was performed by ALFA-SEQ rRNA depletion Kit (Arraystar, USA), and whole mRNAseq libraries were generated using the NEB Next® UltraTM Nondirectional RNA Library Prep Kit for Illumina® (New England Biolabs, MA), following the manufacturer’s recommendations. The index-coded samples were clustered on a cBot Cluster Generation System and sequenced on an Illumina Novaseq6000 platform, generating 150-bp paired-end reads (Guangdong Magigene Biotechnology Co., Ltd., Guangzhou, China). All raw sequences were deposited at the National Center for Biotechnology Information (NCBI) Sequence Read Archive under the accession number PRJNA1011033.

**Figure 1.**
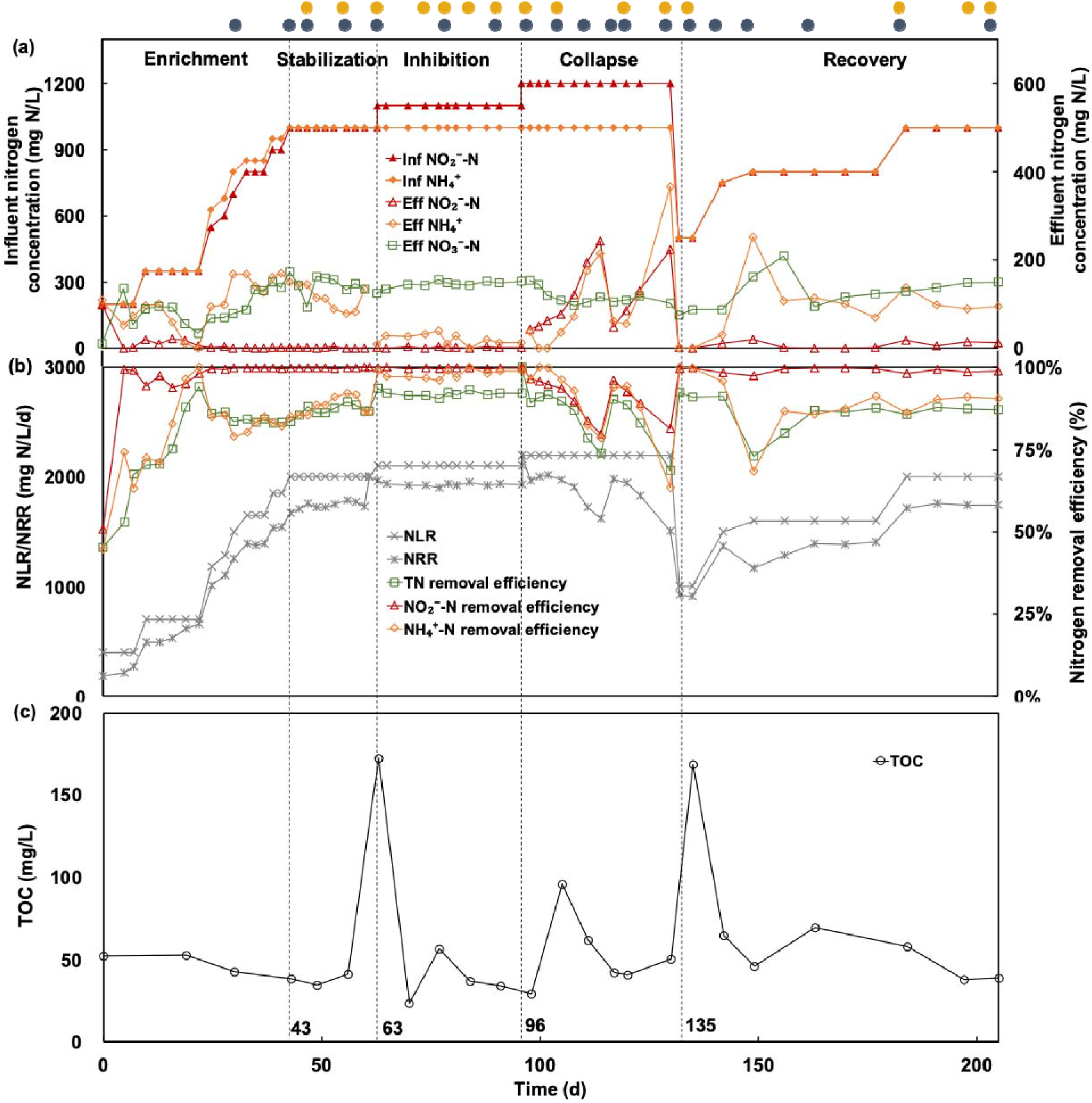
Performance of anammox membrane bioreactor. (a) influent ammonium (NH_4_ ^+^-N) and nitrite (NO_2_ ^-^-N) concentration and effluent ammonium, nitrite, and nitrate (NO_3_ ^-^-N) concentration (mg N/L); (b) removal efficiency (%) of ammonium, nitrite, and total nitrogen (TN), and nitrogen removal rate (NRR) and nitrogen loading rate (NLR) (mg N/L/d). (c) total organic carbon (TOC) concentration (mg/L). Grey and orange dots above x-axis indicated sampling time points for metagenomics and metatranscriptomic analysis.

### 2.5. Metagenomic analyses

Metagenomic analysis was conducted based on our previous study ^6^. Briefly, clean reads were assembled using MEGAHIT version 1.2.2 ^29^. The filtered contigs were binned with BASALT (Binning Across a Series of AssembLies Toolkit) ^30^ to obtain metagenome-assembled genomes (MAGs). MAGs meeting the criteria of completeness – 5 * contamination ≥ 50% were selected for further analysis to infer potential metabolic functionalities ^31^. And then MAGs were assigned taxonomic classifications and calculated relative abundance ^32,33^. Open reading frames (ORFs) were predicted from the MAGs using Prodigal version 2.6.2 ^34^. Predicted amino-acid sequences were annotated against the Kyoto Encyclopedia of Genes and Genomes (KEGG) database through the BLASTP program with an E-value cutoff of 10^-5^. Metabolic pathways were constructed using the KEGG Mapper.

Trimmomatic v. 0.36 was employed to acquire clean data from the raw data of transcriptomics ^35^. Clean reads were then aligned to the NCBI SILVA databases, and rRNA sequences were removed using SortMeRNA ^36^. The remaining mRNA sequences were mapped back onto the MAGs, and the expression level of each gene was quantified using the Reads Per Kilobase Million (RPKM). In order to exclude bias in the comparison of gene expression levels stemming from differences in cell growth, the number of mapped reads for a certain ORF was normalized against the total number of mapped reads for all ORFs within a given genome for RPKM value calculation:

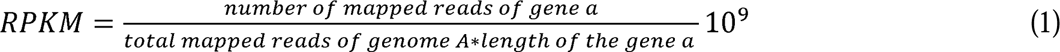

where gene a is present in genome A ^37^. The normalized RPKM value allows a more accurate comparison of gene expression levels within the same genome, rather than across the entire community. To identify differentially expressed genes (DEGs), T-test and Fold Change analysis were performed. The resulting p-values were adjusted using the Benjamini-Hochberg method. Subsequently, DESeq2 package in R v.4.0.5 ^38^ was utilized to visualize DEGs via volcano plot with significance defined as *P* < 0.05 and |log_2_Fold Change (LFC)| >1. DEGs were subjected to enrichment analysis using the KEGG databases by the clusterProfiler package in R v.4.0.5 ^39^. This analysis aimed to elucidate the impacts of nitrite on various metabolic pathways, including nitrogen cycling, carbohydrate metabolism, energy metabolism, lipid metabolism, lipopolysaccharide metabolism, amino acid biosynthesis, coenzyme biosynthesis, and vitamin biosynthesis in microorganisms. Finally, the influencing mechanisms of these impacts on bacterial growth and nitrogen removal from the anammox system were further elucidated.

## 3. Results and discussion

### 3.1. Inhibition effects of nitrite on nitrogen removal and biomass in anammox system

As shown in **Figure 1a** and **1b**, during the proliferation period, the anammox activity was increased along with the increasing influent substrate concentration. A stable nitrogen removal performance was observed with TN removal efficiency and nitrogen removal rate (NRR) reaching more than 85% and 1700 mg N/L/d on day 43, respectively. Throughout the stabilization period, the influent NO_2_ ^-^-N and NH_4_ ^+^-N were all kept constant at 1000 mg N/L. The effluent NO_2_ ^-^-N concentration gradually decreased to negligible, and the removal efficiency of NO_2_ ^-^-N, NH_4_ ^+^-N and TN surpassed 99%, 87% and 87%, respectively. Moreover, MLVSS also increased from 0.8 g/L in the initial seed to above 3 g/L. These results suggested that the proliferation and stabilization periods effectively resulted in significant improvements in both biomass concentrations and nitrogen removal efficiency.

However, during the collapse period, the nitrogen removal performance of the MBR gradually deteriorated as nitrite accumulates in the MBR. NRR decreased from over 1900 mg N/L/d to 1400 mg N/L/d and the TN removal efficiency dropped to 69%. When the effluent NO_2_ ^-^-N concentration rapidly accumulated to 243 mg N/L, the disintegration of granular sludge started to happen. This result was in agreement with previous study reported nitrite acted as an inhibitor for anammox consortia, leading to decreased nitrogen removal performance^40^.

Naturally, reducing NLR is an effective means to mitigate the inhibitory effect from nitrite, as previously reported^28,41^. We took the same approach in our study. During the recovery period, the influent NO_2_ ^-^-N and NH_4_ ^+^-N concentrations were adjusted to 500 mg N/L to reduce NLR until the the MBR performance was gradually restored. Subsequently, the TN removal efficiency and NRR reached over 86% and 1700 mg N/L/d by gradutely increase the ammonium and nitrite concentrations.

The concentration of TOC in the anammox system also exhibited changes corresponding to the ramping up effluent nitrite concentrations (**Figure 1c)**. During the stabilization period, the TOC concentration in the effluent remained constant at 50 mg/L. However, it increased to 173 mg/L and 169 mg/L during the initial inhibition and complete collapse periods, respectively. This observation is in consistent with the previous study, in which environmental stress immediately triggers cells to release cellular components such as soluble microbial products, cytochromes, and extracellular polymeric substances, resulting in increased soluble organic matter ^22,23^. These results indicated that excessive nitrite might cause microbial death.

Additionally, TOC content during the recovery period decreased to a similar level comparable to the stabilization period (around 50 mg/L) accompanied by reduced effluent nitrite concentration. These observations suggest that the microbial consortia reverse back to a normal life status when the nitrite inhibition was lifted. Therefore, it can be speculated that the collapse of the anammox system may be attributed to the direct suppression of nitrogen removal activity by excessive nitrite or detrimental effect on microbial growth. To prove the hypothesis, we conducted further investigations at molecular level to fully understand the underlying mechanisms.

### 3.2 Community structure and temporal dynamics

Genome-resolved metagenomic analysis was conducted to characterize the community structure and microbial functions throughout the whole reactor operation period. A total of 82 MAGs met the standard (completeness - 5 * contamination ≥ 50%) ^31^ were recovered with metagenomic binning from 19 samples. These 82 MAGs were further analyzed and taxonomically classified based on phylogenetic analysis by taxonomic marker genes (**Figure 2**). They belonged to 18 different phyla, with Planctomycetes being the dominant phylum. Following Planctomycetes, the phyla in descending order of relative abundance are Chloroflexi, Bacteroidetes, and Proteobacteria. It is worth noting that Chloroflexi, Bacteroidetes, and Proteobacteria have often been identified alongside Planctomycetes in other bioreactor studies^28^. The relative abundance of Planctomycetes reached 53% after the proliferation stage (**Figure 2A**), which contained a large proportion of AnAOB. In the complete collapse period, the relative abundance of Planctomycetes declined rapidly to 31% before bouncing back to 41% in the recovery period. This suggests that Planctomycetes might be sensitive to the concentration of nitrite. Furthermore, the relative abundance of Bacteroidetes, which includes heterotrophic DNRA and/or canonical denitrifying bacteria, also exhibited significant fluctuations. It has been reported that Bacteroidetes is a major phylum of bacteria with wide distribution and diverse lifestyles and are capable of adapting to various environments ^42^. As they are specialized in the decomposition of complex organic materials ^43^, Bacteroidetes could use high levels of TOC during the collapse period to increase their relative abundance from 9% to 23%. As the concentration of TOC decreased during the recovery period, the relative abundance of Bacteroidetes dropped down to 12%. These results are consistent with the changes in relative abundance observed at the phylum level according to 16S rRNA analysis (**Figure S2**).

**Figure 2.**
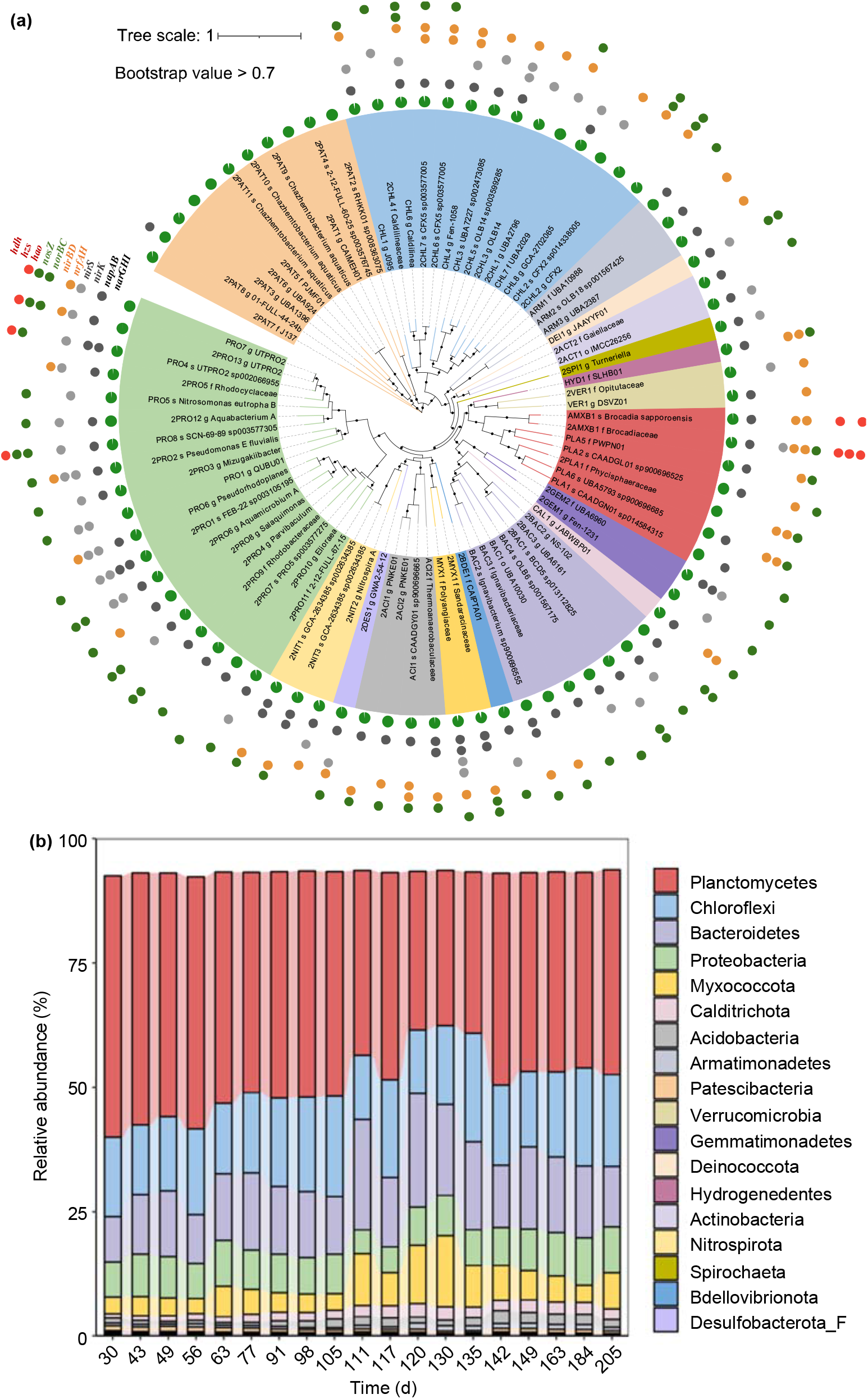
Microbial classification and community structure analysis. (a) Phylogenomic reconstruction of MAGs from anaerobic ammonium oxidation (anammox) bioreactor communities. A phylogenetic tree was generated by maximum likelihood (ML) (inferred with IQ-tree and LG + F + R7 model) with 1,000 bootstrap replications. Different colored tree branches indicated different phylum levels. Bootstrap values > 70% were highlighted with black circles. The green pie represented the completeness of MAGs. Genes in various metabolic pathways were marked with different colored circles by iTOL version 4.4.2. Consequently, pathway of high-quality MAGs was summarized as follows: AnAOB-MAGs found with anammox genes (*hao*/*hzsABC/hdh*), dissimilatory nitrate reduction (DNRA) bacteria-MAGs found with DNRA genes (*nirBD*/*nrfAH*), and denitrifying (DN) bacteria-MAGs found with DN genes (*nirS*/*nirK*/*norBC*/*nosZ*). (b) Composition of bacterial community at phylum over lifespan of bioreactor based on MAGs.

A total of seven MAGs were annotated as Planctomycetes, among which AMXJ1 and AMXB1 were identified as AnAOB because *hzs* and *hdh* genes were discoved in their MAGs. The relative abundance of AMXJ1 was less than 1%, so the focus of the analysis was on AMXB1, which was annotated as *Ca.*Brocadia sapporoensis. Additionally, a comparative genomic analysis of AMXB1 was also conducted (**Figure S3**). AMXB1 had a M bp genome with a GC content of 42%. It contained 2803 ORFs, 3 rRNA, 1 mRNA, and 47 tRNA. Based on average nucleotide identity similarity, AMXB1 was found to have the closest evolutionary relationship with *Ca.*Brocadia sp.UTAMX1 (GCA_002050325.1), showing a similarity of 98.5%. The relative abundance of AMXB1 decreased rapidly from 23% to 8% during the complete collapse period before climbing back to 24% in the subsequent recovery period. These changes in relative abundance were consistent with the results obtained from 16S rRNA gene analysis, although the relative abundance was underestimated (**Figure S2**).

Subsequently, the core metabolic model of AMXB1 was constructed **(Figure S4)**. AMXB1 possessed genes for the transport of substrates such as nitrite transport gene *nirC* and *narK*, ammonium transport gene *amt* and nitrate transport gene *narK*. Previous studies have shown that the nitrite reduction genes *nirS* and *nirK*, which convert nitrite to NO, were not present in *Ca. Brocadia*. However, other nitrite reduction processes may exist^19,44^. Consistent with these findings, AMXB1 also lacked *nirS* and *nirK* genes but possessed hydroxylamine oxidoreductase (HAO), which can reduce nitrite to NH_2_OH^44^. The resulting NH_2_OH and NH_4_ ^+^ are then converted to N_2_ via hydrazine synthase (HZS) and hydrazine dehydrogenase (HDH). In addition, AMXB1 was found to have another nitrite conversion pathway, the dissimilatory nitrate reduction to ammonium (DNRA) pathway, which enables it to reduce nitrite to ammonium ^45^. Furthermore, AMXB1 was capable of assimilating ammonia to glutamate, which was used for subsequent amino acid synthesis and carbon cycling^46^. AMXB1 could synthesize 15 amino acids, including arginine, aspartate, cysteine, glutamine, glycine, isoleucine, leucine, lysine, phenylalanine, proline, tryptophan, tyrosine and valine, which may also be the incomplete of the genome. Moreover, AMXB1 could use the reductive acetyl-CoA pathway to simulate CO_2_ as a carbon source and was capable of completing fatty acid synthesis, as well as partial coenzyme and vitamin biosynthesis. The ability to transport lipopolysaccharides, lipoproteins, putrescine, phospholipids, metal ions, and inorganic salt ions ensures normal growth and metabolism.

### 3.2 Enhanced anammox activity of AMXB1 under excessive nitrite exposure

During the complete collapse, the anammox consortia simultaneously down-regulated the major nitrogen removal genes, including those involved in the anammox pathway, ammonia assimilation, and DNRA pathway (**Figure 3a**). Specifically, the RPKM values of *hao*, *hzs*, and *hdh* genes associated with the anammox pathway decreased from 1102, 8013, and 12460 to 149, 2789, and 3197, respectively. The expression of the *nrfAH* gene involved in the DNRA pathway also decreased from 430 to 104 RPKM. Additionally, the expression of the *gdhA* gene involved in ammonia assimilation decreased from 1662 to 877 RPKM. Consistent with the down-regulation of nitrogen removal gene expression within the overall microbial community, the TN removal efficiency rapidly decreased from 91±1% to a minimum of 69%. While during the recovery period, the nitrogen metabolism genes were upregulated, which perfectly coinciding with the improving nitrogen removal efficiency back to 87±1% (**Figure 3c**).

**Figure 3.**
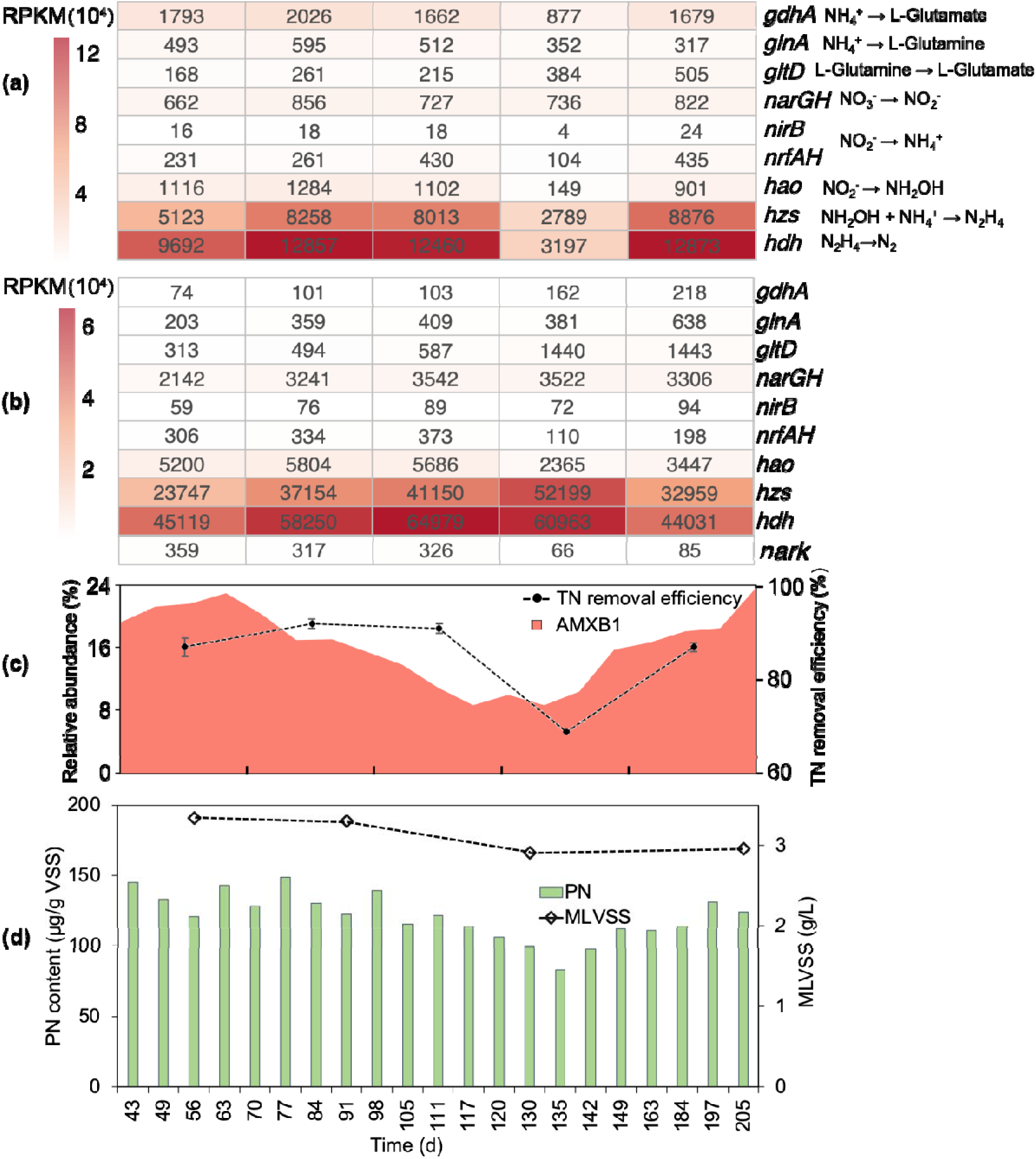
Analysis of the relationship between the expression of nitrogen removal genes and nitrogen removal efficiency of the anammox system. (a) The total nitrogen (TN) removal efficiency and relative abundance of AMXB1. (b) Expression of nitrogen removal genes in anammox community. (c) Expression of nitrogen removal genes in AMXB1. (d) Mixed liquid suspended solids (MLVSS) and total protein concentrations of the anammox system.

Notably, the anammox pathway was the main nitrogen removal process of the microbial consortia residing in the MBR (**Figure 3a**). Thus, we focused more on the AMXB1 for a better understanding of the nitrite inhibition mechanism. To ensure accurate comparisons of gene expression levels and eliminate biases caused by cell growth, we calculated the expression of nitrogen removal genes for AMXB1 alone, rather than the entire community, at different periods (**Figure 3b**). Interestingly, after the normalization, we noticed that the nitrogen removal capacity of AMXB1 was not inhibited by excessive nitrite, but rather the expression of *hzs* and *hdh* genes was significantly up-regulated. Specifically, the expression of *hzs* and *hdh* genes increased from 23747 and 45119 RPKM in the stabilization period to 52199 and 60963 RPKM in the complete collapse period, respectively, which potentially promoted the activity of the anammox process. A similar phenomenon was observed in batch experiments, expression of *hzsB* gene was higher in the presence of 400 mg/L nitrite, which caused inhibition of anammox activity, as compared to non-inhibition conditions ^47^. This is thought to be due to a compensatory mechanism of anammox bacteria for inhibition. In addition, the NarK protein in AnAOB with potential functions of NO_3_ ^-^/NO_2_ ^-^ antiporter can attenuate NO_2_ ^-^ toxicity to anammox cells ^48^. Moreover, energy was required for anammox cells to tolerate NO_2_ ^-^ to drive NO_2_ ^-^ transport ^49^. The *nark* gene was downregulated from 326 to 66 RPKM during the complete collapse period, presumably with less energy being allocated for detoxification. At this point, the excess nitrite could not be pumped out of the cell and could only be rapidly metabolized to prevent more severe cell damage.

However, excess nitrite caused a decrease in the biomass concentration of AMXB1 in the long term. The relative abundance of AMXB1 decreased from 23% to approximately 9% (**Figure 3c**). The MLVSS and total protein concentrations decreased from 3.3 g/L and 136 ± 10 μg/gVSS in the stabilization period to 2.9 g/L and 113 ± 16 μg/gVSS in the complete collapse period, respectively (**Figure 3d**). This suggests that the cells may have diverted less energy for use in reproduction under nitrite stress. During the MBR recovery period, the expression of nitrogen removal genes returned to normal levels and the AMXB1 biomass increased with a decrease in nitrite concentration.

In summary, excessive nitrite greatly reduced the biomass of AMXB1, ultimately resulting in the down-regulation of nitrogen removal gene expression within the entire community and the collapse of MBR performance.

### 3.3 Impaired replication of AMXB1 by excessive nitrite

Previous studies using the amplification of anammox 16S rRNA gene have indicated that nitrite inhibition may be related to microbial growth ^47,50^, but direct evidence is still lacking. To investigate the reasons for the decrease of AMXB1 biomass caused by excessive nitrite, the differentially expressed genes (DEGs) in different periods were analyzed (**Figure 4a**). Then, the DEGs were enriched into the metabolic pathway (**Figure 4b**) and the regulation of microbial functions in different periods was summarized (**Figure 4c**).

**Figure 4.**
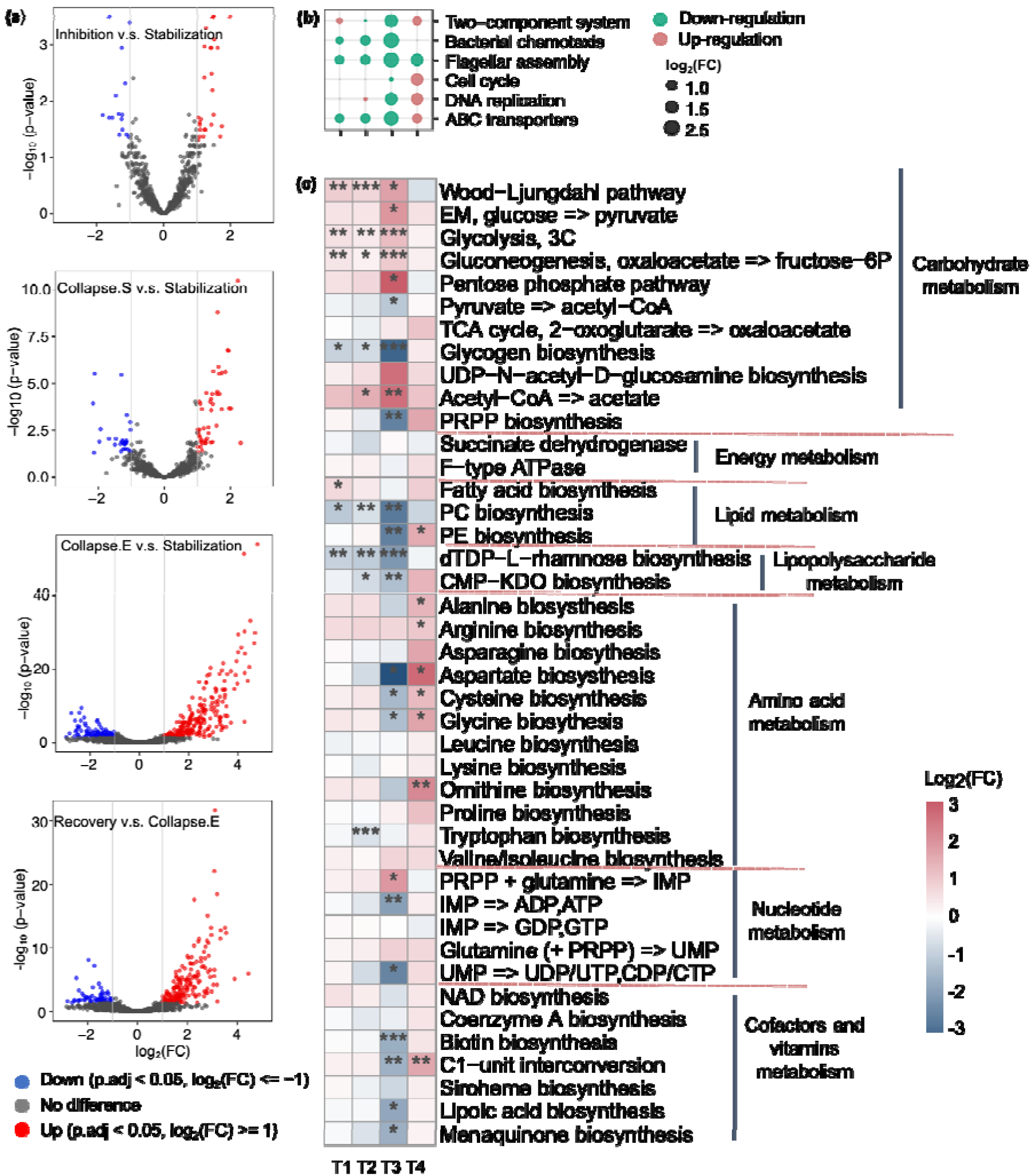
Overview of the effect of nitrite on the core metabolism of AMXB1. (a) The differentially expressed genes (DEGs) of AMXB1 at different periods. (b) Fold change of DEGs for ABC transporters, cell cycle and DNA replication. The size and color of the dots represent log2 transformed fold intensity and directionality. (c) The relative gene transcription associated with core metabolism and biosynthesis of AMXB1. Transcriptomic expression of pathways was relativized by the average RPKM (log2) value of genes in the related pathways. The white box represents the incomplete pathways. (*: *P* < 0.05, **: *P* < 0.01, and ***: *P* < 0.001 by *t*-test). T1: comparison between inhibition and stabilization period, T2: comparison between start of collapse and stabilization period, T3: the comparison between complete collapse and stabilization period, T4: the comparison between recovery and complete collapse period.

It was found that nitrite may inhibit the proliferation of AnAOB by down-regulating certain core metabolic pathways (**Figure 4b**). Firstly, excessive nitrite disrupted the self-protection mechanism of AMXB1. Flagellar assembly and bacterial chemotaxis, which are interdependent processes aiding cells in sensing substance concentration and moving towards nutrients^51,52^, were significantly down-regulated during the complete collapse period. Coupled with the down-regulation of the two-component system that enables bacteria to sense and respond to environmental shock^53^, inferring the weaker sense ability.

Excessive nitrite also prevented AMXB1 from synthesizing important cellular structures (**Figure 4c**). Phospholipids, essential components of the cell membrane ^54^, showed significantly lower synthetic ability during the collapse period compared to the stabilization period. The *malCDE* genes involved in phospholipid intake were down-regulated with 1.4-, 2.4- and 2.3-LFC (*P*-adj. < 0.01), respectively. Lipopolysaccharide, another component of AnAOB cell wall ^55^, exhibited reduced synthesis capacity significantly during the complete collapse but up-regulated during the recovery period. Proteins, which are also components of the cell wall ^56^, showed significant down-regulation of aspartate, cysteine, and glycine during the complete collapse period, indicating a reduction in raw materials for protein synthesis.

Multiple studies collectively identified that excessively high nitrite concentrations may lead to the fragmentation of intact DNA by inducing oxidative stress ^57^. Indeed, excessive nitrites hindered the DNA replication process of AMXB1 (**Figure 4c**). Compared to the stabilization period, cell cycle and DNA replication functions were down-regulated in the complete collapse period. PhosphoRibosyl PyroPhosphate (PRPP), the important intermediate substrate for the synthesis of purine and pyrimidine nucleotides^58^, exhibited reduced synthesis capacity along with other DNA precursors such as adenosine diphosphate (ADP), uridine diphosphate (UDP), and cytidine diphosphate (CDP).

The toxicity of nitrite is expected to disrupt the energy metabolism of AnAOB^48^. C1-unit interconversion and F-type ATP synthesis function, which can provide energy for microorganisms, were down-regulated during the collapse period, indicating insufficient energy supply. However, in this deficient condition, more energy was allocated to carbohydrate metabolism in AMXB1, including glycolysis, the TCA cycle, the pentose phosphate pathway, and Wood-Ljungdahl carbon fixation (**Figure 4c**). Because less energy was devoted to pumping out excess nitrite, the imbalanced energy distribution further impaired self-protection systems, cellular structure generation, and DNA replication.

### 3.4 Transformation of the relationship between AMXB1 and symbiotic bacteria

To associate microbes with different periods of the bioreactor’s lifespan, all MAGs were conducted by pairwise-correlated analysis (**Figure 5a**). The resulting heatmap revealed five distinct clusters (Groups A–E) with AMXB1 included in Group B.

**Figure 5.**
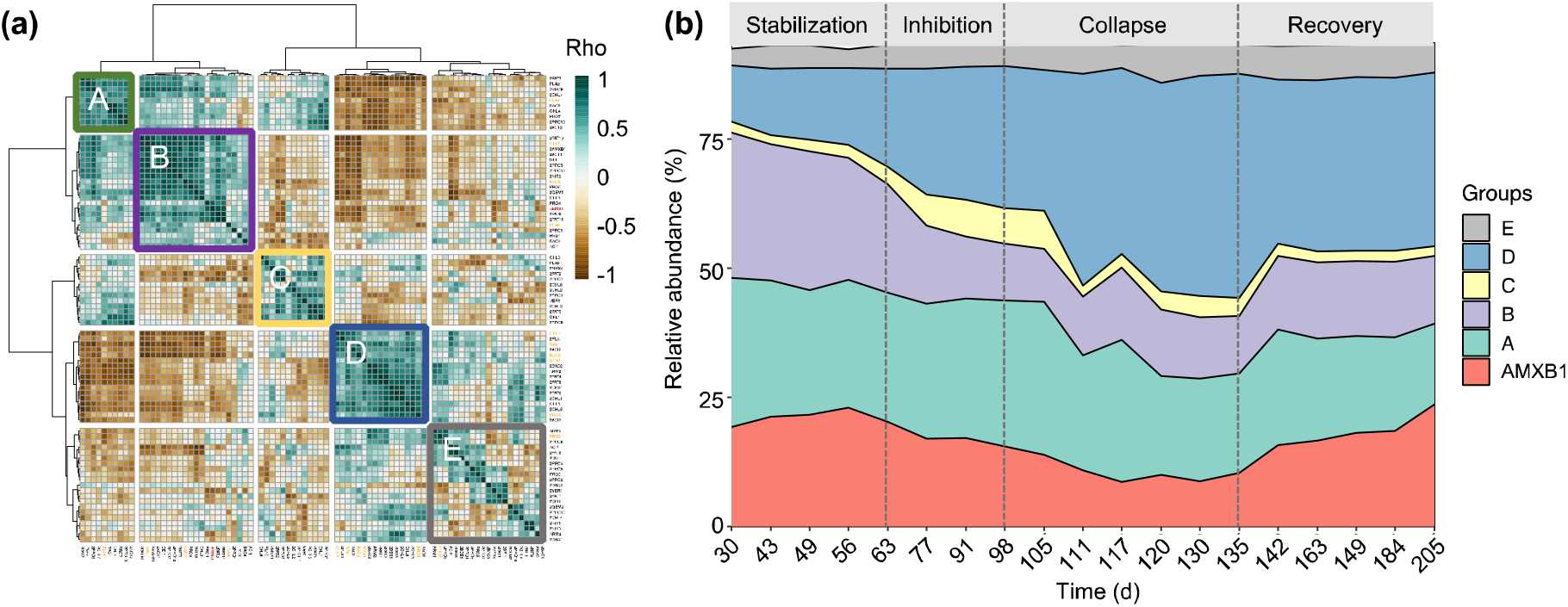
Analysis of the bioreactor’s community clustering, using the relative abundance of the bacteria. (a) A clustering heatmap of bacteria based on pairwise cross-correlations for the 19 time points (matrix values are Rho values). Color scales mark high positive correlation in green and negative correlation in brown. The row and column dendrograms are identical (consisting of the genomes). The row dendrogram shows the calculated distance between the clusters with a dashed red line marking the split into clusters. Colored squares as well as bars to the left of the heat map show relative abundance-based grouping: Green indicates Group A; purple, Group B; yellow, Group C; blue, Group D; and grey, Group E. Red font marks the AnAOB and yellow font marks MAGs > 1%. (b) The relative abundance of different bacterial groups.

The relative abundance of Group A gradually decreased, indicating their association with the initial sludge state, and some of the non-adapted microorganisms disappeared after inoculation. The relative abundance of group E remained relatively constant, suggesting that they were not influenced by nitrite shock.

The relative abundance of Groups C and D increased significantly during the inhibition and collapse periods, with Groups C and D increasing from 3% and 15% in the stabilization period to 4% and 43% in the collapse period, respectively. This could be attributed to the utilization of soluble substances released by autotrophs (either actively or passively) in these groups as carbon sources for proliferation ^22,23^.

The relative abundance variations of these groups were calculated to further examine the role of different microbial groups (**Figure 5b**). The relative abundance of AMXB1 decreased from 23% in the stabilization period to 9% in the collapse period and crepted back in the recovery period. Similarly, the relative abundance variations of Group B were consistent with AMXB1, suggesting a close interaction and symbiotic relationship between them. This indicated that AMXB1 and the microorganisms in group B played a crucial role in maintaining the stability of the anammox system.

Considering that Group B played a crucial role in maintaining the stability of the nitrogen removal performance of MBR, the core metabolic pathways of AMXB1 and major microorganisms in Group B (≥ 1%) were analyzed, including metabolism of carbohydrates, lipids, lipopolysaccharides, amino acids, nucleotides, cofactors, vitamins and energy at different periods. Among them, the functions of BAC2, CHL7 and CHL3 were considerably impacted during the complete collapse and recovery periods. Therefore, a model was established to illustrate the cross-feeding relationship between AMXB1 and these three MAGs by comparing the differential expression of metabolic pathways during nitrite shock and recovery (**Figure 6**).

**Figure 6.**
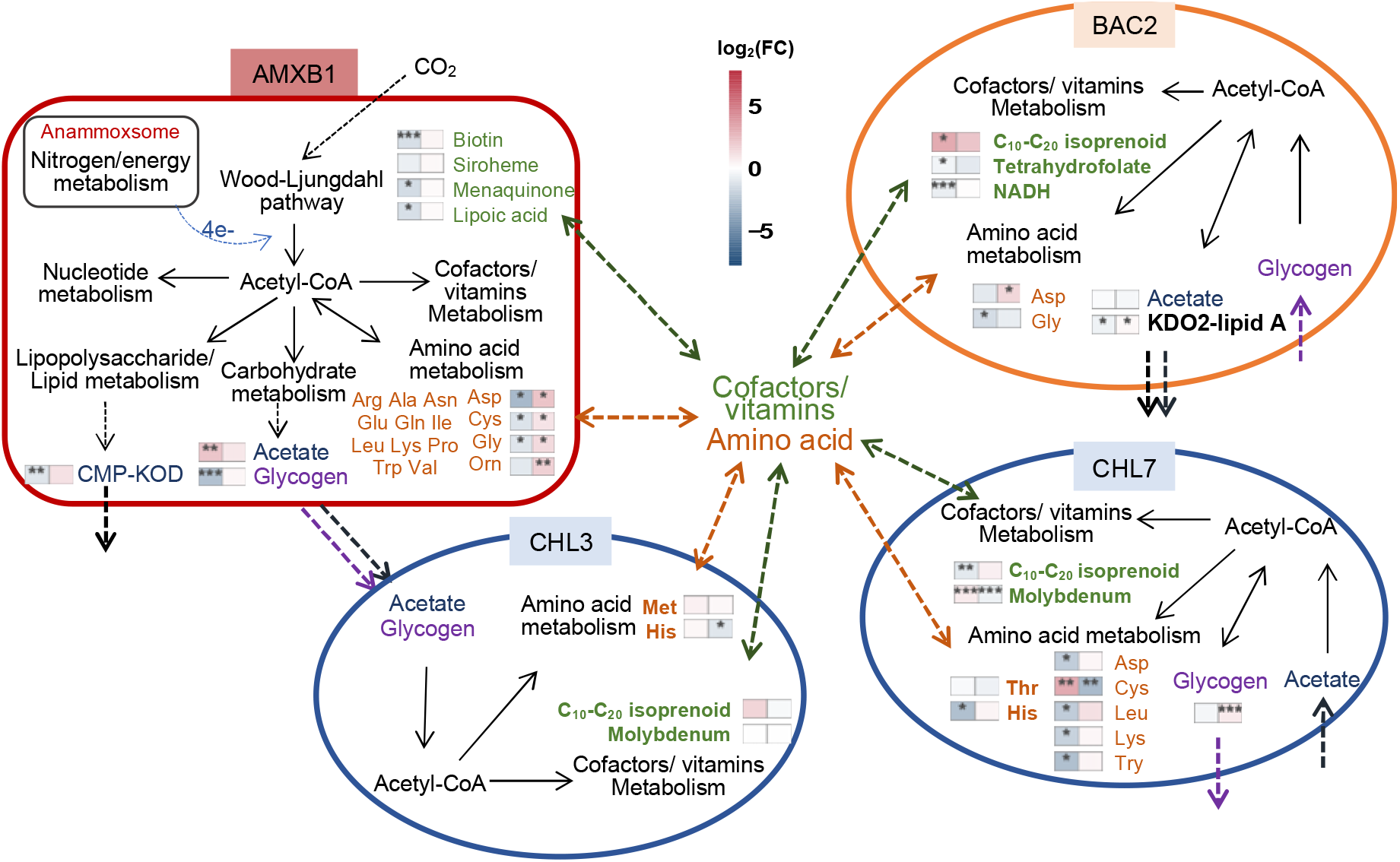
Transformation of the relationship between AMXB1 and symbiotic bacteria. Transcriptomic expression of pathways in the heatmap was relativized by the average RPKM (log2) value of genes in the related pathways. (*: *P* < 0.05, **: *P* < 0.01, and ***: *P* < 0.001 by *t*-test). The square on the left represents the comparison between complete collapse and stabilization period and the square on the left represents the comparison between recovery and complete collapse period. The expression of amino acid synthesis in the figure shows only the amino acids with significant differences and which are lacking in AMXB1 (bold font).

BAC2, CHL7, and CHL3 were capable of nitrogen metabolism through DNRA or canonical denitrification pathways, requiring organic carbons as the electron donor for these processes. However, since there was no organic matter present in the influent, it indicates that microbial cross-feeding is an important means to provide organic substances such as EPS, bacterial cell debris, glycogen, acetate, fatty acids, and more, which was in agreement with the significant increase in TOC concentration (**Figure 1c**). CHL7 and AMXB1 were involved in the glycogen biosynthesis pathway, which could nourish other bacteria within the anammox consortia. Its synthetic capacity was reduced by 3.2-LFC and 2.3-LFC (*P*-adj. < 0.001) during the collapse period and increased by 0.7-LFC (*P*-adj. < 0.001) and 0.3-LFC. Only BAC2 has the ability to synthesize KDO2-lipid A, which is the essential component of lipopolysaccharide in most Gram-negative bacteria and the minimal structural component to sustain bacterial viability ^59^. Its synthetic capacity was reduced by 0.5-LFC (*P*-adj. < 0.01) under nitrite stress but increased by 0.3-LFC (*P*-adj. < 0.01) when inhibition was relieved. Most amino acids were synthesized by AMXB1 to support symbiotic bacteria but were significantly inhibited by excessive nitrite, especially aspartate, cysteine and glycine (**Figure 6**). AMXB1 could not synthesize methionine, histidine and threonine, and the synthesis of histidine was down-regulated 2.9-LFC (*P*-adj. < 0.01) in CHL7 during the collapse period.

AMXB1 was also unable to synthesize folate, a coenzyme that is essential for the growth and activity of AnAOB ^60^, and down-regulated 0.4-LFC (*P*-adj. < 0.01) in BAC2 under nitrite stress. C10-C20 isoprenoid, a component of the bacterial plasma membrane, plays a significant role in electron transport and oxidative phosphorylation ^61^ and depends on CHL7, CHL3 and BAC2 for its production. AMXB1 also could provide most coenzymes to other bacteria (**Figure 6**). Biotin plays a vital role in fatty acid synthesis ^62^. Siroheme is an intermediate in the synthesis of heme in prokaryotes and plays an important role in the reduction of nitrite^63^. Lipoic acid participates in acyl transfer in substance metabolism and eliminates damaging free radicals ^64^. Menaquinone is a component of the electron transport chain in most anaerobic bacteria to produce ATP^65^. These functions experienced significant down-regulation during the complete collapse period and up-regulation during the recovery period. Therefore, the cross-feeding of glycogen, lipopolysaccharide, amino acid and coenzyme between AMXB1 and symbiotic bacteria was hindered by excess nitrite. This hindrance was reversed as the concentration of nitrite decreased. Previous studies collectively indicated the cross-feeding of nutrients within anammox systems has the potential to regulate and influence the activity of AnAOB ^18^. Hence, the close cross-feeding relationship facilitated the exchange of essential substances and played a crucial role in restoring the nitrogen removal performance of the anammox system.

## 4. Conclusions

In this study, we conducted an in-depth investigation into the effects of excessive nitrite concentrations on the core metabolism of AnAOB and symbiotic bacteria within an anammox consortia. The overarching goal was to uncover the mechanism underlying the effects of nitrite on microbial growth and nitrogen removal performance. It was observed that nitrite at 243 mg N/L caused complete collapse of nitrogen removal in a continuous-flow anammox MBR. Metagenomic and metatranscriptomic analyses revealed that excessive nitrite disrupted the energy metabolism of AMXB1 and less energy was devoted to establishing tolerance and cell replication, contributing to the eventual collapse of the anammox system. Additionally, the cross-feeding of glycogen, lipopolysaccharide, and amino acids between AMXB1 and symbiotic bacteria was hindered by excess nitrite. By reducing the nitrite concentration, the microbial consortia gradually regained their nitrogen metabolisms and overall performance. These findings provide insights into the nitrite inhibition mechanisms of AnAOB and highlight the important role of a close cross-feeding relationship in the recovery of nitrogen removal performance within the anammox system.

## Acknowledgments

This work was financially supported by the National Natural Science Foundation of China (NO. 51709005), Shenzhen Knowledge Innovation Program Basic Research Projects (JCYJ20190808183205731 and JCYJ20220812103301001). This work was also supported by the Shenzhen municipal development and reform commission (discipline construction of watershed ecological engineering). W.-Q.Z. thanks the Catalyst: Leaders funding (CHN-UOA1601) provided by the New Zealand Ministry of Business, Innovation and Employment, and administered by the Royal Society Te Apārangi.

