## Supplemental File for "Nitrite deteriorates bioreactor performance by reducing growth of *Ca.* Brocadia sapporoensis instead of inhibiting the anammox activity"

**Document prepared:**

Number of Pages: 11

Number of Texts: 2

Number of Figures: 4

Number of Table: 1

**Text S1. EPS extraction**

10 mL of sludge-water mixture was taken from MBRs and centrifuged at 4,000 rpm for 10 min, the pellets were washed three times with 10 mM phosphate buffered saline at pH 7.5, and then re-suspended to a pre-determined volume. CRE (DOWEX™ MARATHON™ CA) was added as a dosage of 70g/g VSS. The suspensions were magnetically stirred for 2 h at 1200 rpm. After removing the CER, EPS were harvested at 12,000 rpm and 4 °C for 5 min. The supernatants were filtered through 0.45 µm acetate cellulose membranes (Advantec Co., Japan).

**Text S2. Metagenomic analyses**

For shotgun metagenomic sequences, raw paired-end reads were initially filtered using fastp ^1^. Paired-end reads were filtered when the number of low-quality bases (Q ≤ 5) exceeded 50% in any sequence read. Filtered reads were assembled using MEGAHIT version 1.2.2^2^ specifying a range of k-mer size values (79-149) in 20 base pair intervals and finally reserved contigs > 1,000 base pairs.

The filtered contigs were processed with BASALT (Binning Across a Series of AssembLies Toolkit) ^3^ to obtain bins. The completeness and contamination of the bins were then estimated using CheckM version 1.0.13 ^4^ with lineage-specific marker genes and default parameters. Bins that meet the MIMAG standard (completeness -5*contamination ≥ 50%) ^5^ were retained in this study as metagenome-assembled genomes (MAGs) to infer potential nitrogen metabolism functionalities.

120 bacteria-specific protein sequences of conserved maker-genes were identified from MAGs using GTDB-Tk version 0.3.2 ^6^. Each protein sequence was individually aligned using hmmalign ^7^ with default parameters. Individual alignments were concatenated and used as input to reconstruct the phylogenomic tree using the IQ-TREE version 1.6.12 ^8^ with the best-fit model at 1,000 times bootstrapping (-b 1000). Finally, the phylogenetic tree was visualized and edited in the iTOL version 4.4.2 (https://itol.embl.de/personal_page.cgi) online platform ^9^.

The relative abundances of the MAGs were calculated by several steps. The MAGs were firstly combined as a reference file and bowtie2 ^10^ was used to construct a comparison template, that is, to establish an index. Next, each set of raw data was mapped to the reference file, then a customized script was used for coverage and length calculations (<http://github.com/banfieldlab/mattolm-public-scripts>). The relative abundance of MAGs was calculated by the average coverage multiplying the length of each MAG divided by the total read base pairs in each sample ^11^ as shown in Eq. (S1).

$MAG abundance=\frac{coverage * length of MAG}{total reads of each sample}$	(S1)

Genes were predicted from the MAGs using Prodigal version 2.6.2 ^12^, and predicted amino-acid sequences were annotated against the Kyoto Encyclopedia of Genes and Genomes (KEGG) database via the BLASTP program against NCBI nr database with an E-value cutoff of 10^-5^. Nitrogen metabolic pathways were constructed using the KEGG Mapper (http://www.genome.jp/kegg/tool/map_pathway.html) to visualize results.

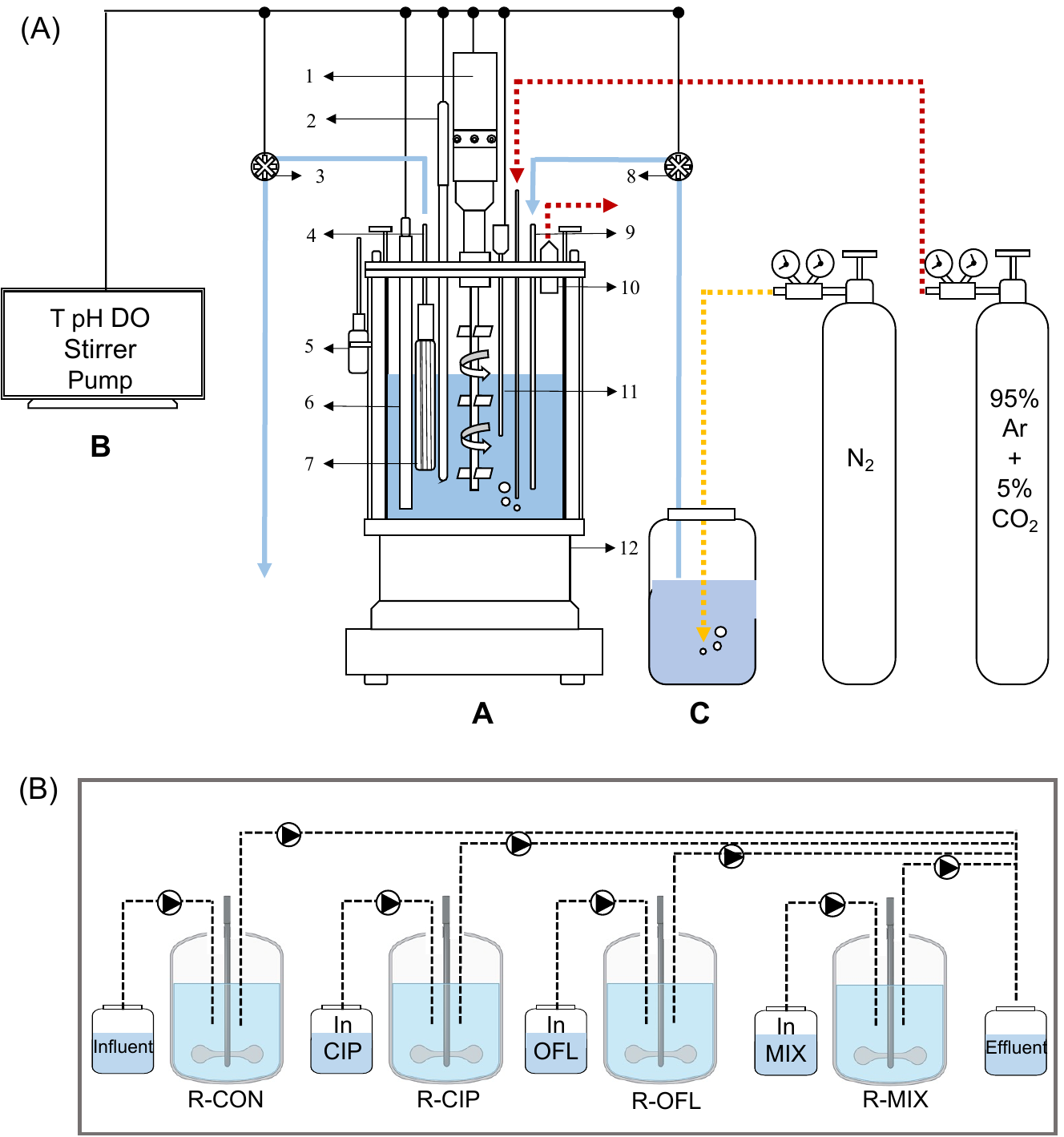

**Figure S1.** The structure of membrane bioreactor (MBR) system. A – MBR; B – control panels; C – medium with brown bottles. 1 – stirrer; 2 – temperature sensor; 3 – effluent pump; 4 – effluent; 5 – sample collection bottle; 6 – sampling tube; 7 – membrane; 8 – inffluent pump; 9 – influent; 10 – exhaust unit; 11 – DO sensor; 12 – water bath unit.

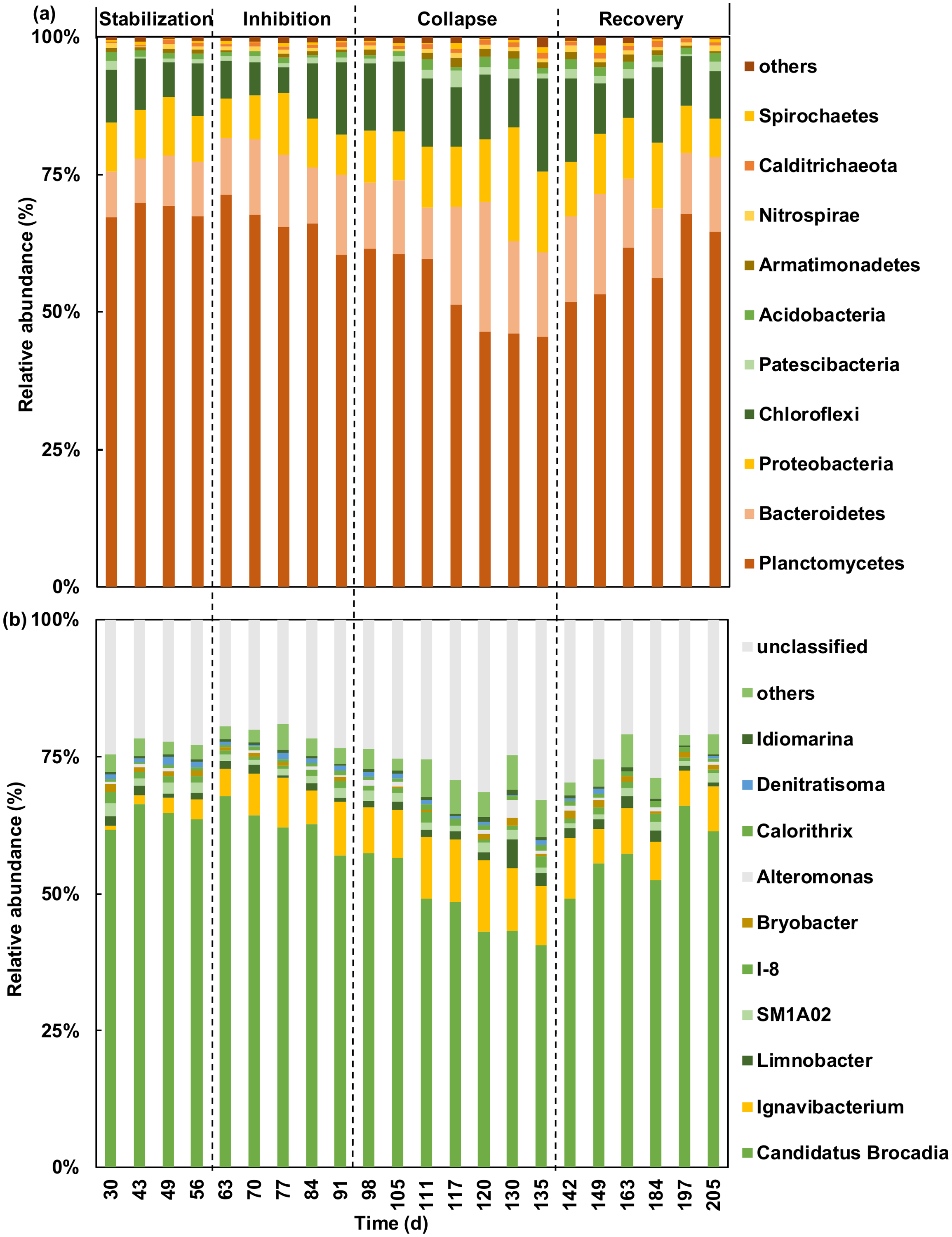

**Figure S2.** Composition of bacterial community at phylum (A) and genus (B) levels in samples over lifespan of bioreactor based on 16S rRNA OTUs.

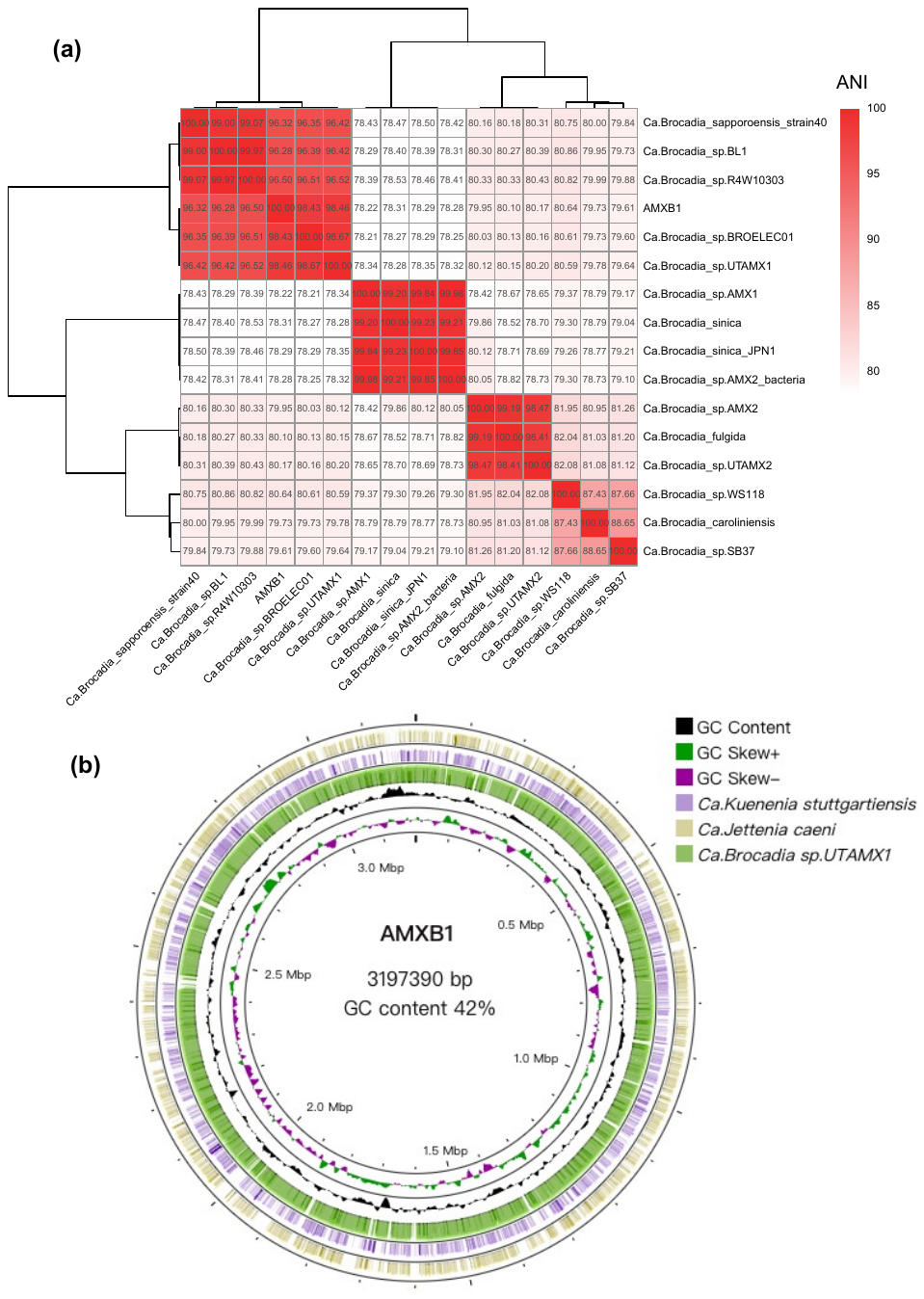

**Figure S3.** Comparative genomic analysis of AMXB1. (a) The average nucleotide similarity (ANI) heatmap of AMXB1 and AnAOB MAGs downloaded from NCBI. (b) Difference analysis between AMXB1 and other AnAOB MAGs.

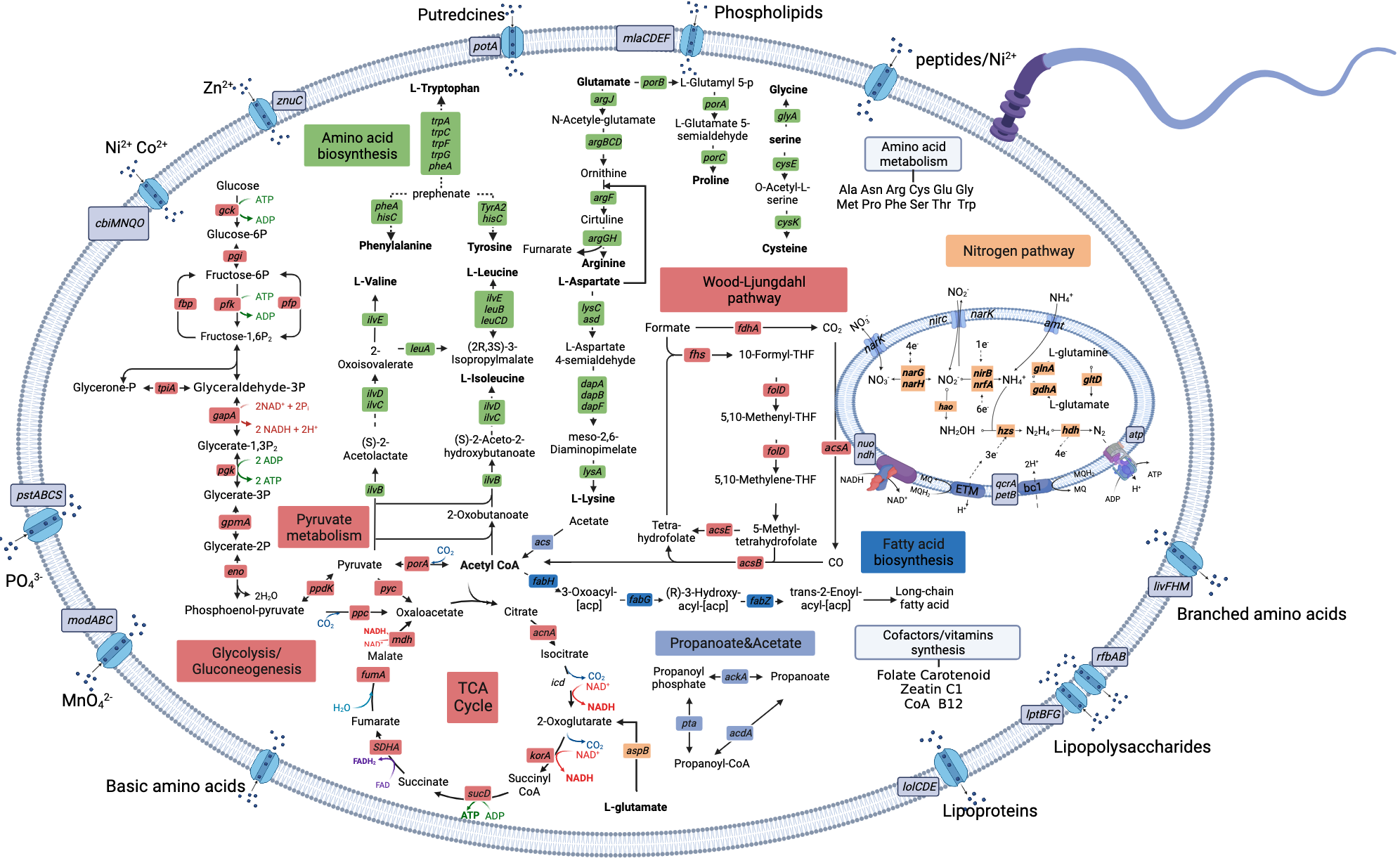

**Figure S4**. Core metabolic model of AMXB1

**Table S1.** **The time points for taking biological samples**

| Times (days) | Total protein | 16S rRNA analysis | Metagenomic analysis | Metatranscriptomic analysis |
| --- | --- | --- | --- | --- |
| 30 |  | √ | √ |  |
| 43 | √ | √ | √ |  |
| 49 | √ | √ | √ | √ |
| 56 | √ | √ | √ | √ |
| 63 | √ | √ | √ | √ |
| 70 | √ | √ |  | √ |
| 77 | √ | √ | √ | √ |
| 84 | √ | √ |  | √ |
| 91 | √ | √ | √ | √ |
| 98 | √ | √ | √ | √ |
| 105 | √ | √ | √ | √ |
| 111 | √ | √ | √ |  |
| 117 | √ | √ | √ |  |
| 120 | √ | √ | √ | √ |
| 130 | √ | √ | √ | √ |
| 135 | √ | √ | √ | √ |
| 142 | √ | √ | √ |  |
| 149 | √ | √ |  |  |
| 163 | √ | √ | √ |  |
| 184 | √ | √ | √ | √ |
| 197 | √ | √ | √ | √ |
| 205 | √ | √ | √ | √ |
